# VizR: An Interactive Web Platform for End-to-End RNA-Seq Analysis and Visualization in Plant Biology

**DOI:** 10.64898/2026.07.24.740467

**Authors:** Woo-Taek Jeon, Hoon Jung, Donghwan Shim, Yuree Lee

## Abstract

RNA sequencing (RNA-seq) is widely used to investigate transcriptional programs in plant biology, yet the need to combine multiple specialized tools and bioinformatics expertise to convert raw sequencing reads into biologically interpretable results remains a major technical barrier for many plant biologists. Here, we present VizR (VIsualiZation of Rna seq), a web-based platform that integrates end-to-end RNA-seq analysis and visualization within a single integrated environment. VizR automates upstream processing, including quality control, adapter trimming, genome alignment, and transcript quantification, and connects the resulting expression data to downstream exploratory analyses. Its interface is designed to make expression patterns immediately searchable and interpretable: users can query genes through an equalizer-style expression-pattern interface, inspect expression profiles using inline heatmaps embedded in gene tables, and perform context-integrated gene ontology analysis throughout the workflow. VizR also supports comparative analysis through interactive Venn diagram module, allowing users to transfer gene sets directly from result tables. As a Docker-based application, VizR can be deployed locally and accessed through a standard web browser. By unifying automated RNA-seq processing, interactive visualization, and functional interpretation, VizR lowers the technical barrier to transcriptome analysis and provides a practical platform for plant biology research.

## Introduction

Next-generation sequencing (NGS) technologies have established RNA sequencing (RNA-seq) as a routine and powerful approach for transcriptome profiling in plant biology (Wang et al., 2009). By capturing genome-wide changes in gene expression, RNA-seq has become central to studies of plant development, physiology, and environmental responses, providing insight into the regulatory networks that shape plant growth and adaptation (Shin et al., 2025; Tsai et al., 2025; Jeon et al., 2026a; Jeon et al., 2026b). In parallel, the rapid accumulation of publicly available plant RNA-seq datasets in repositories such as the NCBI Sequence Read Archive (SRA) has created new opportunities for comparative and integrative transcriptome analysis (Leinonen et al., 2010).

Despite its broad adoption, RNA-seq analysis remains technically challenging for many plant biologists. Although the experimental generation of RNA-seq data has become increasingly routine, the computational workflow required to transform raw reads into biologically interpretable results remains fragmented. Researchers typically need to combine command-line tools for read processing and genome alignment, statistical frameworks for differential expression analysis, and separate software platforms for functional enrichment or pathway interpretation. Because these steps often rely on different file formats, software environments, and levels of computational expertise, each transition can introduce an additional barrier to analysis (Conesa et al., 2016; Pola-Sánchez et al., 2024). As a result, RNA-seq analysis can impose a substantial burden on research groups without dedicated bioinformatics support and delay the extraction of biological meaning from transcriptome datasets (Upton et al., 2023).

The challenge becomes even more pronounced after differential expression analysis, when researchers must translate statistical outputs into biological hypotheses. At this stage, they often need to identify genes with specific expression patterns, compare gene sets across experimental contrasts, and connect transcriptional changes to functional annotations. These exploratory tasks frequently require customized scripts or additional external tools, limiting the ability to move seamlessly from result tables to iterative biological interpretation. Several web-based platforms have been developed to reduce these barriers. Galaxy provides a graphical environment for constructing modular bioinformatics workflows (Afgan et al., 2018), whereas tools such as iDEP enable interactive differential expression analysis and pathway visualization from pre-processed count matrices (Ge et al., 2018). Although these platforms have substantially improved the accessibility of RNA-seq analysis, important gaps remain. Many user-friendly tools begin with count matrices and therefore require upstream processing to be completed separately. Moreover, downstream outputs are often handled as static result tables rather than reusable gene sets that can move seamlessly between visualization, comparison, and functional interpretation modules. These limitations become especially problematic in plant transcriptomic studies involving multiple genotypes, treatment conditions, or dense temporal sampling, where biological interpretation depends on the ability to iteratively explore complex expression patterns.

To address these limitations, we developed VizR, an interactive web platform that integrates upstream RNA-seq processing, downstream visualization, gene-set comparison, and functional interpretation within a single browser-based environment. VizR automates key processing steps from data acquisition to alignment and quantification, and connects the resulting expression data to interactive tools for exploratory analysis. To demonstrate its utility in a realistic plant transcriptomics context, we applied VizR to a publicly available *Arabidopsis thaliana* leaf RNA-seq dataset examining transcriptional responses to mechanical wounding (Lee et al., 2025). This dataset includes 36 samples, with three biological replicates collected across twelve time points, providing a temporally resolved test case for evaluating VizR’s capacity for integrated transcriptome exploration. Through this analysis, we show that VizR enables users to move seamlessly from raw data processing to differential expression analysis, temporal pattern discovery, gene-set comparison, functional annotation, and pathway-level interpretation within a single accessible environment.

## Materials and Methods

### System architecture and implementation

The VizR is freely available at GitHub (https://github.com/snupcb2018/VizR). VizR was implemented as a Docker-based containerized web application to enable reproducible deployment while minimizing system-specific configuration requirements. The server-side application was developed in Python 3.11 using Flask 3.0 and was responsible for analytical workflow execution, user authentication, data storage, and communication with client applications. The client-side interface was implemented using React 19 and TypeScript, compiled with Vite, and connected to the backend through a RESTful API. User session information and workbench metadata were stored in a SQLite database. Real-time updates on pipeline execution status were delivered to connected clients via WebSocket communication using Flask-SocketIO, with Redis serving as the message broker. To support concurrent access by multiple users, VizR generated an isolated directory structure for each user, with separate directories assigned for uploaded files, intermediate files, and analysis outputs. Workbench- level results could be shared with other registered users in read-only mode, enabling collaborative inspection of analysis results while preventing modification of the original workbench.

### RNA-seq processing pipeline

The upstream RNA-seq processing pipeline was implemented as a series of sequential modules for read quality assessment, read preprocessing, genome alignment, transcript quantification, and differential expression analysis (Fig. 1). Each processing step was managed by a multi-worker job scheduler, enabling parallel execution across multiple samples. Raw sequencing reads were initially evaluated using FastQC (Andrews, 2017). Adapter sequences and low-quality bases were removed using Trimmomatic 0.39 (Bolger et al., 2014). The trimmed reads were further processed with PRINSEQ to remove reads with low sequence complexity or excessive duplication (Schmieder and Edwards, 2011). The resulting high-quality reads were aligned to the reference genome using either HISAT2 (Kim et al., 2019) or Bowtie2 (Langmead and Salzberg, 2012), according to the user-defined pipeline configuration. Alignment files were converted to sorted BAM files using SAMtools (Danecek et al., 2021). Gene-level expression was quantified using StringTie (Pertea et al., 2015), which generated TPM, FPKM, and raw count matrices. Differential expression analysis was performed using edgeR (Robinson et al., 2010) within an isolated R-based Docker container. The edgeR workflow applied trimmed mean of M-values (TMM) normalization followed by negative binomial model-based statistical testing to identify differentially expressed genes.

**Fig. 1.**
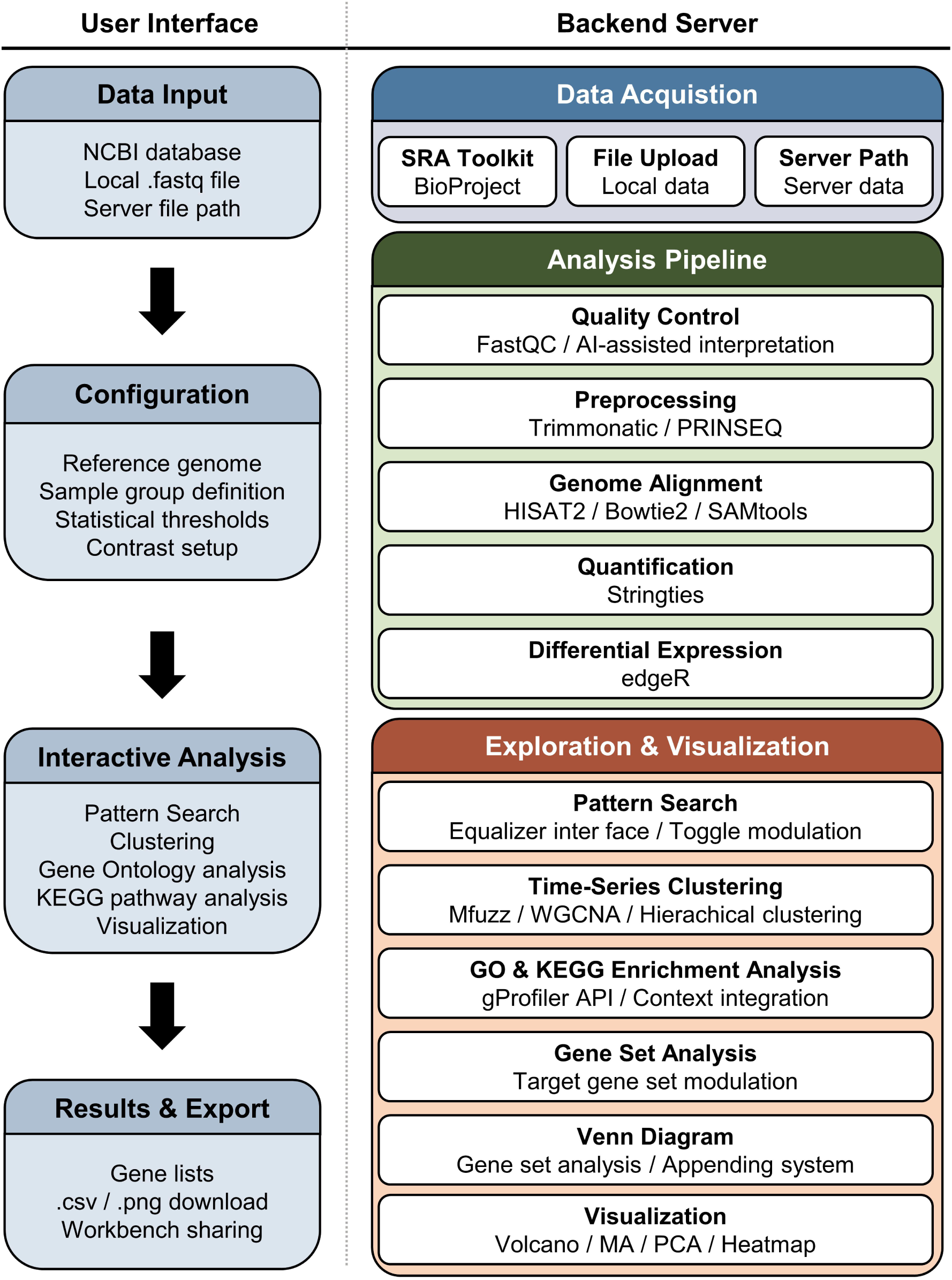
Architecture of the VizR RNA-seq analysis platform. VizR is composed of a web- based user interface and a backend server that together support end-to-end RNA-seq analysis. The user interface enables data input through BioProject accession retrieval, NCBI data upload, or server file path specification, and provides experimental configuration functions for sample group assignment, library layout detection, and pipeline setup. It also integrates downstream analysis modules for gene-pattern search, Venn diagram analysis, clustering, and GO/KEGG enrichment, together with visualization tools for heatmap generation and curated gene set display. Processed results can be exported for further analysis and reporting. The backend server manages data acquisition, executes the upstream RNA-seq workflow, and supports downstream exploration, visualization, and result management.

### Workbench creation and sample configuration

VizR provides a guided workbench creation workflow for RNA-seq analysis starting from raw sequencing data. During workbench setup, users specified the workbench name, target species, and data source. Three input routes were supported: local FASTQ file upload through a drag-and-drop interface, direct retrieval of sequencing data from NCBI using a BioProject accession, and linking of sequencing files already available on the server. For NCBI-based input, VizR automatically retrieved SRA metadata and allowed users to select samples for download. After data selection, sequencing files were assigned to experimental groups and biological replicates through a sample-mapping interface. Paired-end and single-end sequencing layouts were detected automatically. Users then configured the analysis pipeline by selecting the reference genome, alignment method, quality-control parameters, and preprocessing options. Upon completion of workbench setup, VizR initiated the configured RNA-seq pipeline and linked the sample configuration to downstream analysis modules.

To demonstrate the analytical workflow of VizR, the publicly available *Arabidopsis thaliana* RNA-seq dataset GSE275743 was analyzed. This dataset was deposited in NCBI GEO under BioProject PRJNA1152896 and was associated with a study of wound-induced barrier formation in mature leaves (Lee et al., 2025). The dataset comprised 36 samples, consisting of three biological replicates collected across 12 post-wounding time points from 0 h to 5 d after wounding. Raw FASTQ files were retrieved from the NCBI Sequence Read Archive using the VizR data acquisition interface. Reads were aligned to the *A. thaliana* TAIR10 reference genome (Lamesch et al., 2012) using HISAT2 (Kim et al., 2019). Gene-level expression was quantified using StringTie (Pertea et al., 2015), and differential expression analysis between post-wounding time points was performed using edgeR (Robinson et al., 2010).

### Differentially expressed genes analysis

Differentially expressed genes (DEGs) were identified from the edgeR output using a fold-change threshold of >2 and a statistical significance cutoff of *p* < 0.05 (Robinson et al., 2010). For heatmap visualization, expression values of selected genes were transformed into Z-scores across samples on a gene-by-gene basis. Z-score heatmaps were generated in R using the pheatmap package v1.0.12 (Kolde and Kolde, 2015). Gene clustering was performed using the Ptree method implemented in Trinity (Haas et al., 2013), with the Ptree parameter set to 40.

### Visualization modules

VizR included multiple interactive visualization modules to facilitate exploratory analysis of RNA-seq datasets. These modules supported principal component analysis (PCA), volcano plots, MA plots, and replicate scatter plots. Interactive plots were rendered using Plotly and Highcharts, enabling users to inspect sample relationships, expression distributions, and differential expression patterns directly within the web interface. An Interesting Gene Sets module was also implemented to allow users to define, store, and revisit named gene lists. For each stored gene set, VizR generated full-sample Z-score heatmaps, enabling rapid inspection of selected gene families, marker genes, or functionally related gene groups across all experimental conditions.

### Expression matrix visualization and inline mini-heatmaps

Gene expression tables, including DEG tables, expression matrices, clustering outputs, and Venn diagram-derived gene lists, were displayed with an inline mini-heatmap for each gene. Mini-heatmaps were rendered as SVG elements and generated by applying Z-score normalization across samples for each gene. The normalized values were mapped to a user-defined three-color gradient representing low, intermediate, and high expression levels. For DEG-derived tables, the stroke color of each mini-heatmap indicated the direction of differential expression, with red denoting upregulated genes and purple denoting downregulated genes. Hover interaction was implemented for each heatmap cell, allowing users to inspect the corresponding sample name, raw expression value, and Z-score. This design enabled users to evaluate gene-level expression patterns directly within tabular outputs without opening separate visualization panels.

### WGCNA analysis

A signed gene co-expression network was constructed using the WGCNA package v1.72 in R (Langfelder and Horvath, 2008). Prior to network construction, genes were filtered according to expression variance, and the top 5,000 most variable genes were retained for analysis. Pairwise correlations between genes were calculated across samples, and the resulting correlation matrix was transformed into a signed adjacency matrix using the formula 0.5 + 0.5 × correlation. The adjacency values were then raised to a soft-thresholding power, which was automatically selected by the pipeline. Network construction was performed using the following parameters: input source type, variance-filtered genes; top variable genes, 5,000; soft-thresholding power, auto-detected; minimum module size, 100; deepSplit, 4; and merge cut height, 0.1. The adjacency matrix was converted into a topological overlap matrix (TOM), which provided a robust measure of network interconnectedness. Genes were then grouped by average-linkage hierarchical clustering based on TOM-derived dissimilarity. Co-expression modules were identified using the Dynamic Tree Cut algorithm applied to the resulting hierarchical clustering dendrogram. Gene lists from each co-expression module were subjected to Gene Ontology enrichment analysis using the g:Profiler API (Kolberg et al., 2023).

### Pattern-based gene discovery using equalizer

Pattern-based gene discovery was implemented through an equalizer-style graphical interface. In this module, users defined an expected expression pattern by adjusting vertical sliders corresponding to experimental sample groups. The backend calculated the Spearman rank correlation coefficient between each gene’s group-median expression profile and the user-defined pattern vector. Genes were retained when the correlation coefficient met or exceeded the user-defined minimum Spearman threshold. Additional filtering options were implemented to refine pattern-based gene searches. These included a minimum log₂ fold-change between the highest and lowest expression groups, minimum CPM, maximum coefficient of variation, and equality or ordering constraints between selected group pairs. Individual sample groups could be toggled on or off, allowing users to include or exclude specific groups from the pattern-matching constraint. Output genes were ranked in descending order according to Spearman correlation, enabling prioritization of genes whose expression profiles most closely matched the user-defined pattern.

## Results

### End-to-End Processing of a Public Plant RNA-seq Dataset Using VizR

To demonstrate the complete analytical workflow supported by VizR, we processed the publicly available RNA-seq dataset PRJNA1152896, which consists of 36 samples collected across twelve time points after mechanical wounding. Starting from the BioProject accession in the data acquisition interface, VizR automatically retrieved raw FASTQ files from NCBI SRA and detected the paired-end sequencing layout without manual specification (Fig. 2A). The full pipeline, including FastQC quality assessment, Trimmomatic/PRINSEQ preprocessing, HISAT2 alignment to the TAIR10 reference genome, StringTie quantification, and edgeR differential expression analysis, was executed within a single workbench session (Fig. 2B). This workflow was managed by VizR’s multi-worker job scheduler, allowing parallel processing across samples.

**Fig. 2.**
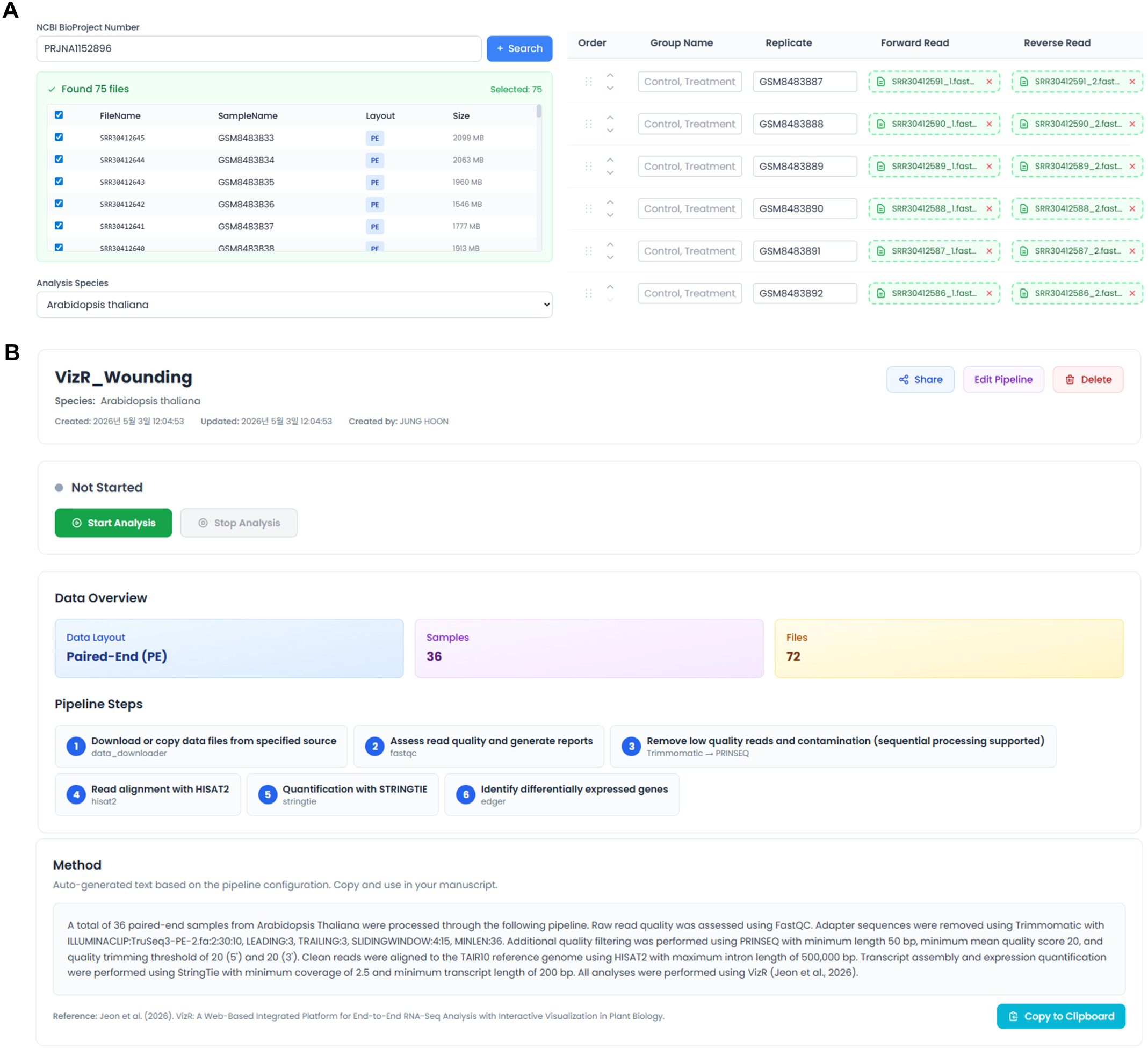
Workbench creation interface. VizR provides a guided workbench creation interface that integrates BioProject-based data retrieval, sample annotation, and pipeline configuration for reproducible RNA-seq analysis. Users initiate a workbench by entering a BioProject accession number, after which VizR retrieves associated metadata from NCBI and supports selective download of SRA files. Retrieved or uploaded files are then organized into experimental groups and biological replicates through a sample-mapping interface, with automatic detection of paired-end and single-end library layouts. Before execution, users specify the reference genome, quality control tool, and preprocessing parameters, thereby generating an analysis-ready workbench for downstream RNA-seq processing.

The dataset showed consistently high sequencing quality across all samples. FastQC analysis indicated a mean per-base Phred quality score of 36.3, and the QC Analysis module displayed the six core FastQC metrics across selected samples, enabling rapid identification of potential quality outliers without requiring users to inspect individual reports separately (Fig. 3). Following adapter trimming and low-complexity filtering, reads aligned efficiently to the TAIR10 genome, with uniformly high alignment rates across samples (99.3–99.5%) (Fig. 4A– C). StringTie quantification generated expression estimates for 32,833 genes, corresponding to the total annotated gene set of *Arabidopsis thaliana*.

**Fig. 3.**
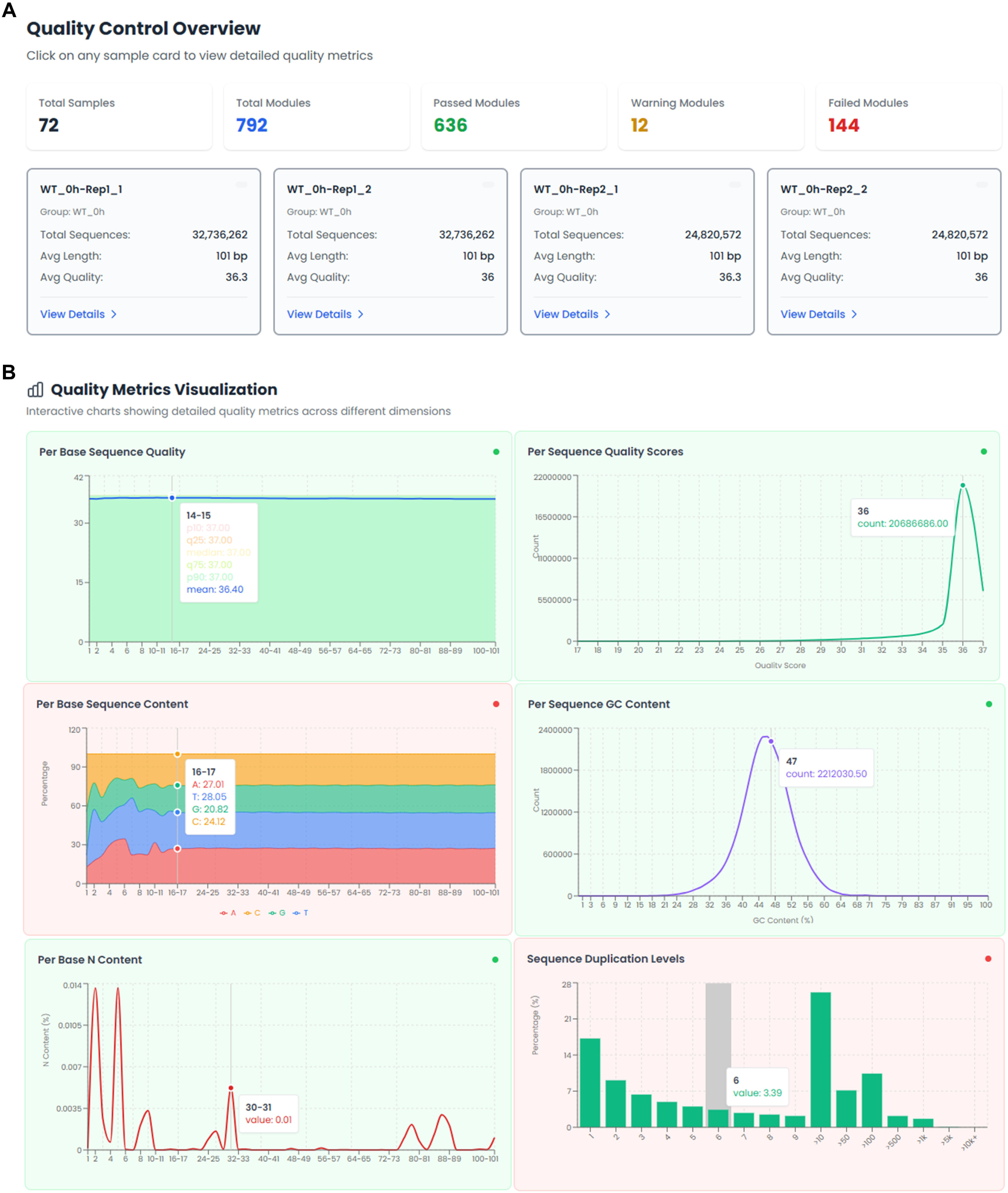
Quality control results. (A) Quality Control Overview displaying FastQC-derived quality metrics across all samples. Sample-specific cards report the total number of sequences, average read length, and overall quality scores, with color coding indicating pass, warning, or fail status. (B) Representative quality metric visualization for WT_0h-Rep1-1, showing detailed FastQC-derived profiles used to assess read quality and sequencing consistency prior to downstream analysis.

**Fig. 4.**
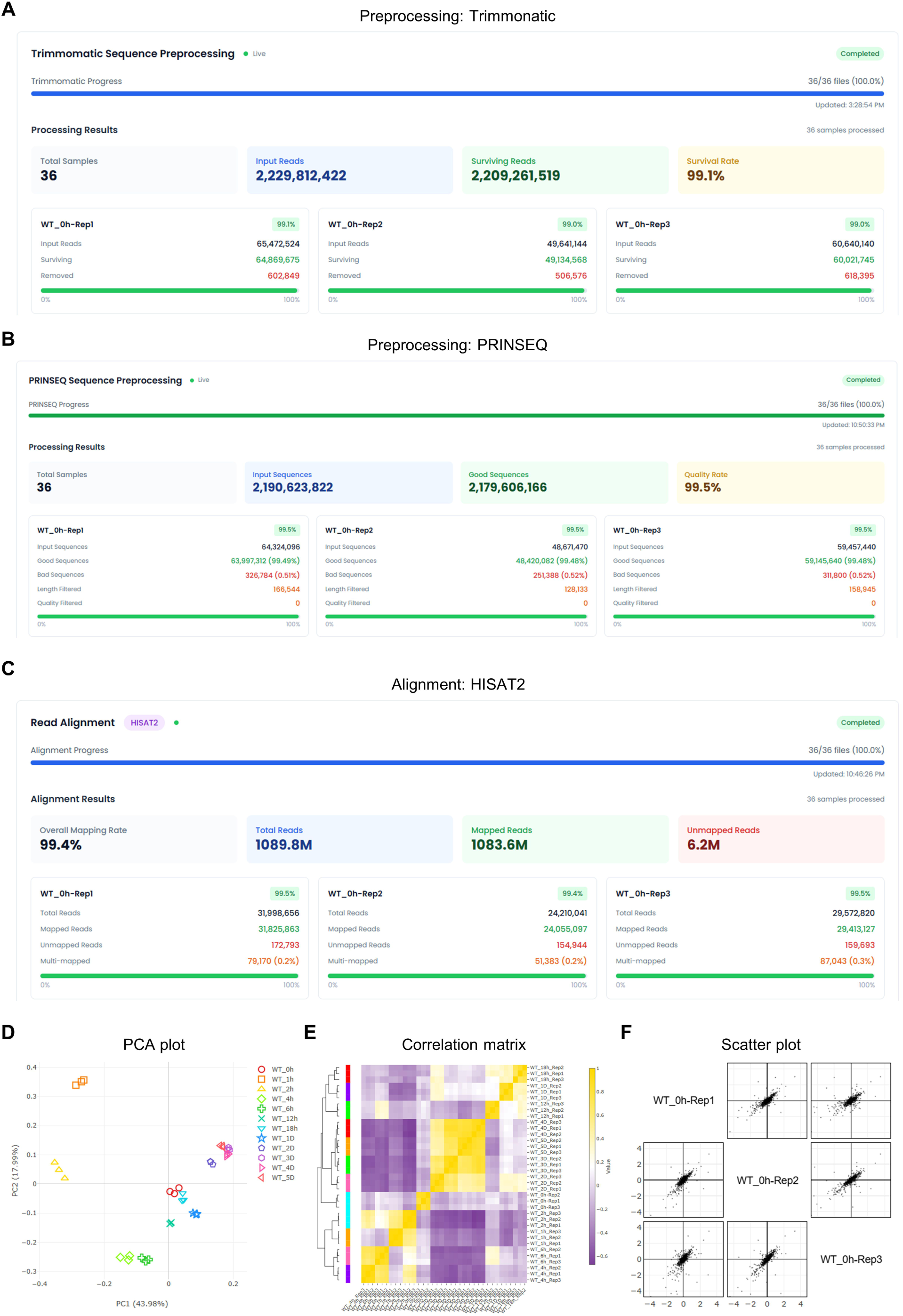
Preprocessing and genome alignment results. (A) Trimmomatic preprocessing results showing adapter trimming and quality filtering across 36 samples, with 2,229 M input reads, 2,209 M surviving reads, and an overall survival rate of 99.1%. (B) PRINSEQ preprocessing results showing additional quality filtering, with 2,190 M input reads, 2,179 M surviving reads, and an overall survival rate of 99.5%. (C) HISAT2 genome alignment results showing an overall mapping rate of 99.4%, with per-sample summaries of total, mapped, and unmapped reads. Across all 36 processed samples, 1,083 M reads were mapped out of 1,089 M total reads, whereas 6 M reads remained unmapped. (D) Principal component analysis (PCA) of the full expression matrix showing separation of time points along PC1 and PC2, which explained 43.98% and 17.99% of the total variance, respectively. Biological replicates clustered closely within each condition, indicating high reproducibility. (E) Correlation matrix showing expression similarity among sample groups. (F) Pairwise replicate comparison within the WT_0h group, showing concordant expression patterns among biological replicates.

Differential expression analysis was performed using edgeR for 66 contrasts defined through the VizR group configuration interface. Using thresholds of |log₂FC| > 1 and FDR < 0.05, we identified 14,925 differentially expressed gene calls across all contrasts. The strongest transcriptional difference was observed between 2 hours and 4 days post-wounding, yielding 7,815 DEGs, including 3,688 genes upregulated at 2 hours and 4,127 genes upregulated at 4 days post-wounding. PCA of the full expression matrix confirmed the reproducibility of the dataset, with biological replicates clustering tightly and samples separating according to time point along the first two principal components (PC1: 43.98%, PC2: 17.99%) (Fig. 4D). Consistent with this pattern, the correlation matrix revealed distinct relationships among time points, while replicate-level scatter plots showed strong concordance among biological replicates within each time point (Fig. 4E, F). Together, these results demonstrate that VizR can execute a complete RNA-seq workflow from public data acquisition to differential expression analysis, while producing quality-controlled and reproducible outputs suitable for downstream temporal transcriptome exploration.

### Interactive Reuse of Gene Sets Across Analysis Modules

Beyond executing the upstream and differential expression workflow, VizR treats result tables as interactive entry points for downstream analysis. Genes selected from DEG result tables, expression tables, clustering results, or Venn diagram regions can be transferred directly to heatmap visualization, GO enrichment, KEGG pathway analysis, or comparative gene-set analysis without file export (Fig. 5A). DEG result tables also include inline mini heatmaps that allow users to inspect expression profiles while filtering and selecting candidate genes (Fig. 5B). This design enables users to move iteratively between statistical results, expression-pattern inspection, functional interpretation, and gene-set comparison within a single analytical environment.

**Fig. 5.**
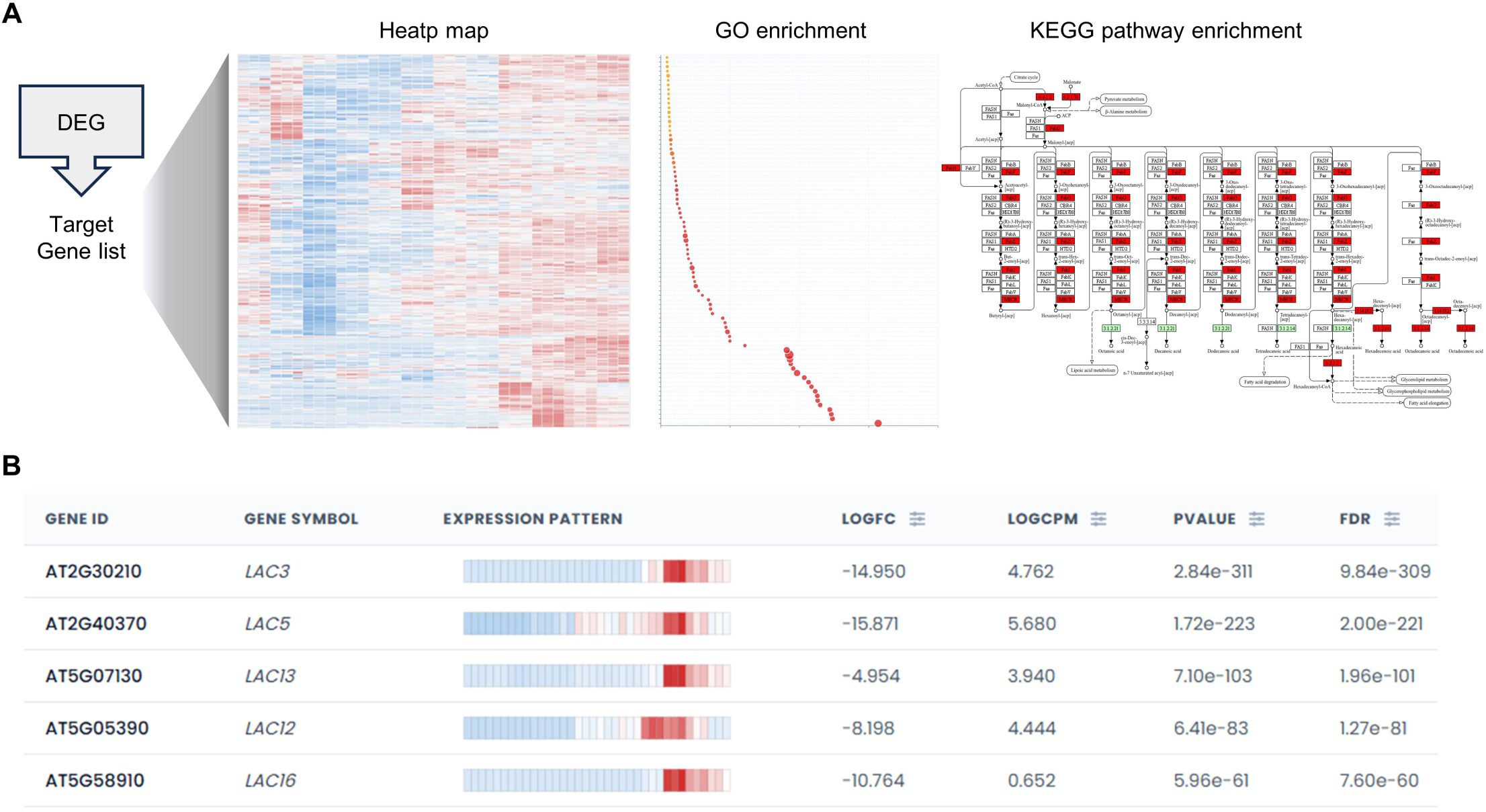
Interactive Gene-Set Reuse in VizR. (A) Schematic overview of the interactive gene- set reuse framework in VizR. Genes selected from DEG tables can be directly transferred to downstream modules, including heatmap visualization, GO enrichment, and KEGG pathway analysis, without manual export or reformatting. (B) Representative DEG table with embedded inline mini heatmaps, enabling rapid inspection of expression patterns during candidate gene filtering and downstream interpretation.

### Interactive Clustering and Functional Annotation of Wound-Responsive Gene Sets

After differential expression analysis, a major challenge is to organize large DEG lists into biologically interpretable groups with shared expression dynamics. To support this step, VizR provides multiple clustering approaches for expression-profile analysis. Hierarchical clustering with user-selectable linkage methods, including Ward, average, and complete linkage, organizes genes into expression-based groups and displays the results together with an interactive dendrogram. Users can adjust the dendrogram cut point to define the desired number of clusters, and the mean expression pattern of each cluster is rendered as a line chart, enabling rapid visual comparison across clusters (Fig. 6A).

**Fig. 6.**
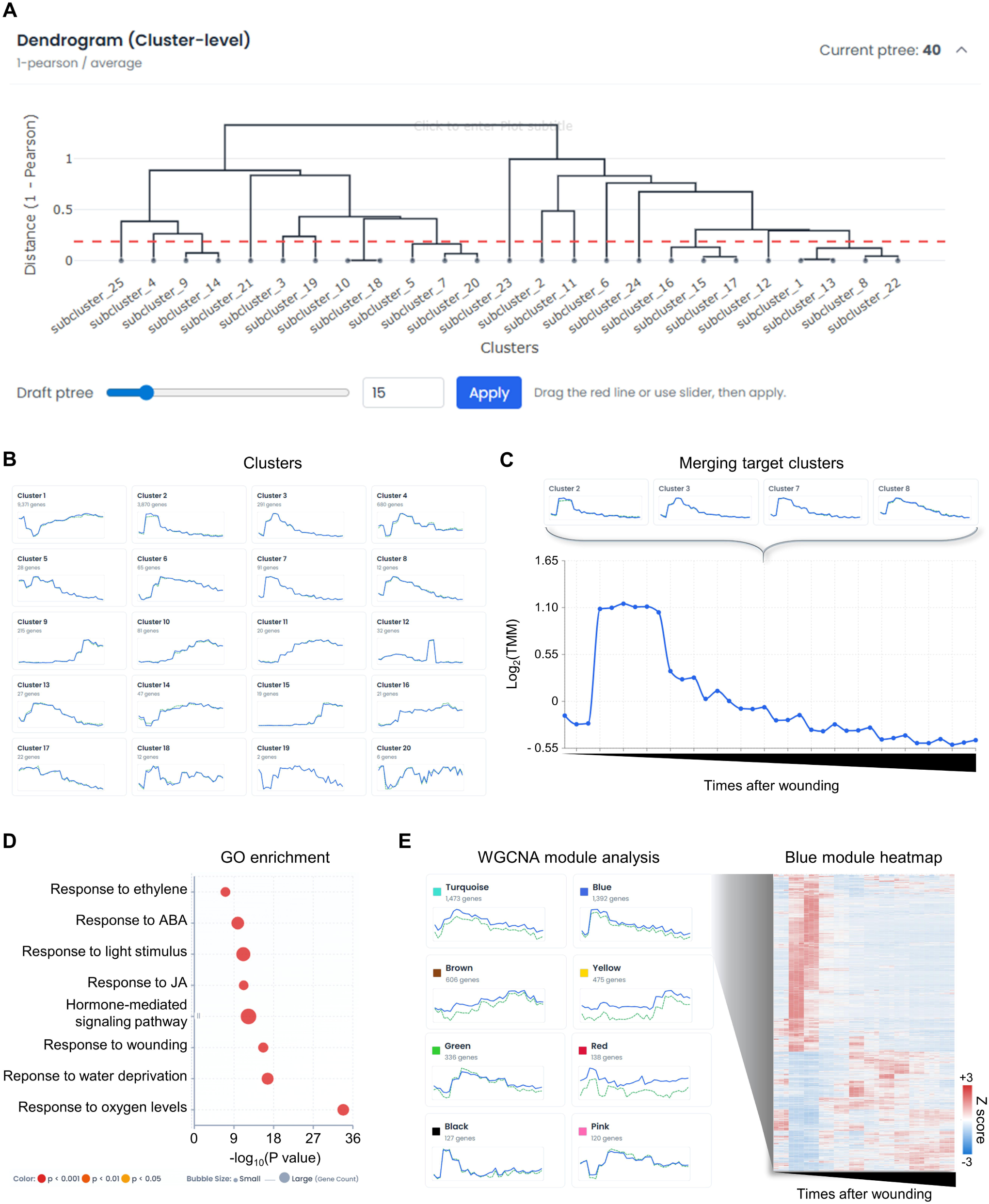
Cluster analysis results. (A) Interactive interface for controlling cluster number by adjusting the Ptree value and dendrogram cut point. (B) Twenty expression clusters generated from differentially expressed genes (DEGs) using edgeR-based expression pattern analysis, revealing distinct temporal transcriptional profiles across the dataset. (C) VizR enables user-defined merging of clusters with selected expression patterns. In this example, clusters showing increased expression at early time points after wounding were combined into a single merged cluster for focused downstream analysis. (D) Gene Ontology (GO) enrichment analysis of the merged cluster, showing biological processes enriched among genes associated with the early wound-induced transcriptional response. (E) Co-expression-based module analysis of the full gene set using WGCNA. A heatmap of the Blue module is shown as a representative module, illustrating coordinated expression patterns among genes assigned to this co-expression group.

To demonstrate how VizR supports user-guided gene set curation, we applied hierarchical clustering to differentially expressed genes and partitioned them into 20 expression clusters (Fig. 6B). We then focused on clusters associated with early transcriptional responses to wounding. Four clusters showing induction between 1 h and 4 h after wounding, Clusters 2, 3, 7, and 8, were merged within VizR and directly subjected to Gene Ontology (GO) enrichment analysis (Fig. 6C). The merged gene set was enriched for GO terms associated with hormone responses, including abscisic acid, ethylene, and jasmonic acid, which have been implicated in early wound signaling (Fig. 6D). The same gene set was also enriched for terms related to environmental responses, including water- and oxygen-associated processes (Fig. 6D). These results indicate that early wound-induced transcriptional reprogramming is associated with both hormonal and environmental response pathways.

VizR also supports weighted gene co-expression network analysis (WGCNA), which is particularly useful for temporal datasets or complex experimental designs. In this analysis, WGCNA separated genes into co-expression modules based on shared expression patterns and identified the blue module as an early wound-responsive module (Fig. 6E). The blue module showed elevated expression during the early post-wounding period, supporting its association with early wound-responsive transcriptional programs. Together, these clustering and module-based analyses demonstrate that VizR can extract biologically meaningful temporal gene groups from large RNA-seq datasets and connect these groups directly to functional enrichment analysis.

### Pattern-Guided Gene Discovery Using the Equalizer Interface

Clustering-based analysis is useful for identifying previously unrecognized expression patterns in an unbiased manner. However, in many RNA-seq studies, the biological process of interest is already suggested by prior phenotypic or physiological observations, and the analytical goal is to identify genes that follow a specific expected trajectory. In such cases, repeatedly adjusting clustering parameters to recover a desired pattern can be inefficient and may obscure biologically relevant gene sets. To support hypothesis-guided gene discovery, VizR provides an equalizer-style pattern search interface that allows users to define an expected expression profile directly by adjusting per-group sliders, without requiring prior knowledge of gene identities (Fig. 7A).

**Fig. 7.**
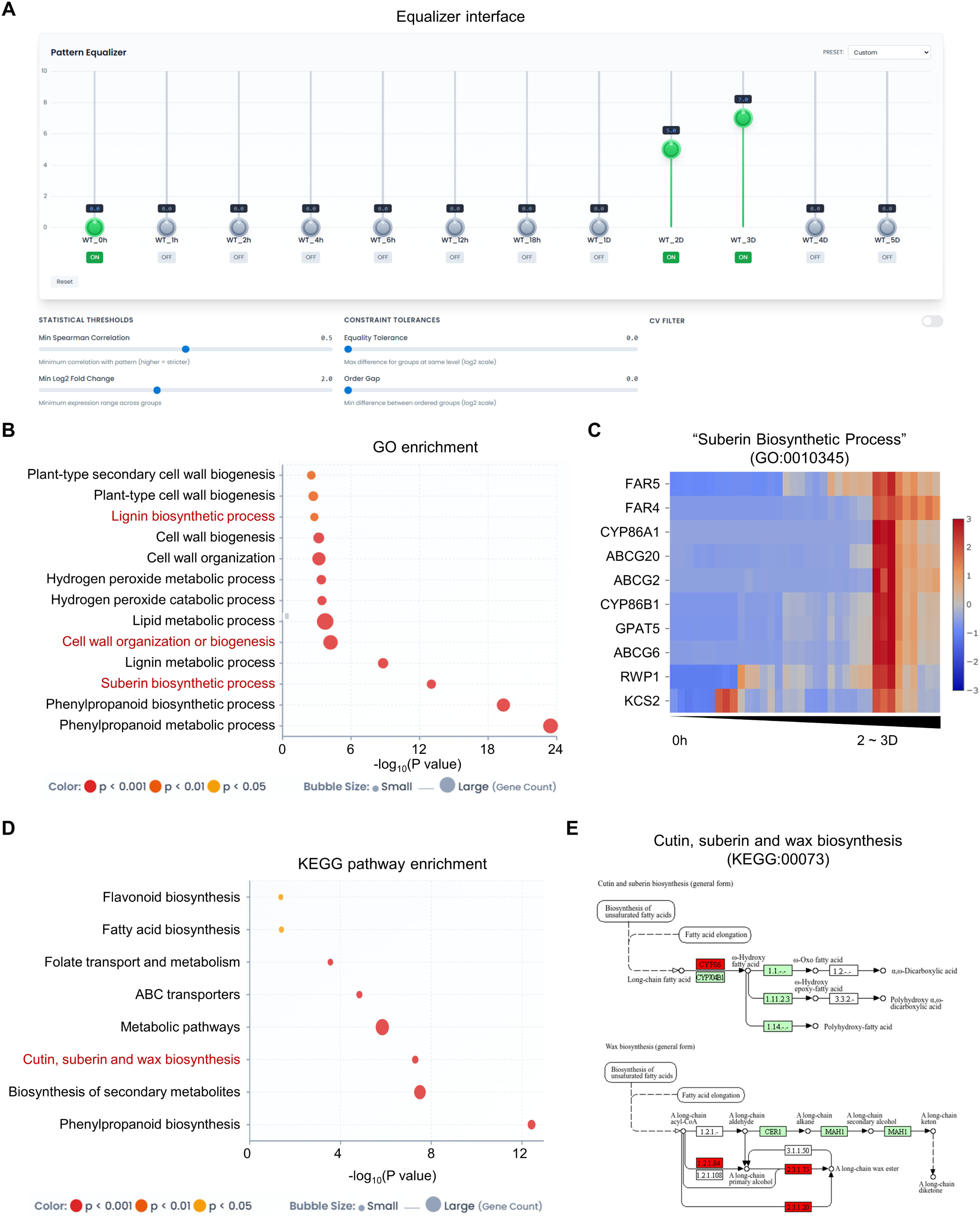
Equalizer-based pattern search and functional enrichment analysis. (A) Equalizer- style pattern search interface used to identify genes showing sustained late wound induction. (B) Gene Ontology (GO) enrichment analysis of the 541 pattern-matched genes, showing enrichment of suberin biosynthesis, cell wall organization or biogenesis, and lignin biosynthesis. (C) Expression profiles of genes associated with the enriched GO term “suberin biosynthetic process,” including representative suberin-related genes such as GPAT5, FARs, and ABCGs. (D) KEGG pathway enrichment analysis of the same pattern-matched gene set, highlighting enriched pathways including fatty acid biosynthesis, ABC transporters, and cutin, suberin, and wax biosynthesis. (E) Pathway-specific expression patterns of genes assigned to the enriched KEGG pathway “Cutin, Suberin and Wax Biosynthesis.”

To illustrate this capability, we searched for genes showing sustained late wound induction, characterized by low expression at the unwounded baseline, gradual induction by 2 days post-wounding, and peak expression at 3 days post-wounding. This trajectory is consistent with the temporal progression of secondary barrier formation described by Lee et al. (2025) (Lee et al., 2025), in which suberin and lignin deposition become apparent during the later phase of wound healing. The desired expression pattern was defined by positioning three representative time-point sliders, WT_0h, WT_2D, and WT_3D, at low, intermediate, and high values, respectively, while intermediate time points were toggled off to exclude them from the pattern constraint (Fig. 7A). By simultaneously applying a minimum Spearman correlation threshold of ρ ≥ 0.5 and a minimum log₂ fold-change threshold of ≥2, VizR retrieved 541 genes, which were ranked in descending order according to their correlation coefficients.

GO enrichment analysis was then performed directly within VizR without file export or switching to external tools. Biological process enrichment analysis identified 29 significantly enriched terms among the 541 pattern-matched genes. Prominent enriched terms included “suberin biosynthetic process” (GO:0010345; 11 genes; *p* = 9.40 × 10⁻¹⁴), “cell wall organization or biogenesis” (GO:0071554; 33 genes; *p* = 6.33 × 10⁻⁵), and “lignin biosynthetic process” (GO:0009809; eight genes; *p* = 1.65 × 10⁻³) (Fig. 7B). Selecting the enriched term “suberin biosynthetic process” enabled direct inspection of the corresponding genes and their expression profiles within VizR (Fig. 7C). This gene set included canonical components of suberin biosynthesis and transport, such as *GLYCEROL-3-PHOSPHATE sn-2- ACYLTRANSFERASE 5* (*GPAT5*), *FATTY ACID REDUCTASEs* (*FARs*), and *ATP-binding cassette subfamily G transporters* (*ABCGs*). The coordinated induction of these genes suggests that the biosynthetic and transport machinery associated with suberin formation is activated during wound-induced secondary barrier formation

The same pattern-matched gene set was further subjected to KEGG pathway enrichment. Eight pathways were significantly enriched, including “Fatty acid biosynthesis,” “ABC transporters,” and “Cutin, suberin and wax biosynthesis” (Fig. 7D, E). These KEGG results were consistent with the GO enrichment analysis and further supported the enrichment of lipid metabolic and transport pathways associated with secondary barrier formation. Together, these results demonstrate that the equalizer-based search successfully recovered genes associated with late wound-induced secondary barrier formation. Thus, VizR enables hypothesis-guided gene discovery by allowing users to translate an expected biological trajectory into a direct transcriptome-wide search, complementing unbiased clustering-based approaches.

## Discussion

RNA-seq analysis remains technically fragmented, requiring users to move across separate tools for data acquisition, preprocessing, alignment, quantification, differential expression analysis, visualization, and functional interpretation. VizR was developed to reduce this barrier by integrating these steps into a single web-based workbench. Rather than requiring users to assemble independent command-line tools and downstream analysis platforms, VizR provides a continuous analytical environment in which public RNA-seq datasets can be retrieved, processed, analyzed, and explored through an interactive browser interface.

A central design principle of VizR is that intermediate results are not treated as static outputs, but as reusable entry points for further analysis. In conventional RNA-seq workflows, DEG tables, expression matrices, clustered gene groups, and comparative gene sets are often exported as separate files and re-entered into different tools. This process can interrupt biological interpretation, particularly when users need to inspect expression profiles, compare gene sets, or connect candidate genes to functional annotations. VizR addresses this problem by allowing genes selected from DEG result tables, expression tables, clustering results, or Venn diagram regions to be transferred directly to heatmap visualization, GO enrichment, KEGG pathway analysis, or comparative gene-set analysis. In this way, VizR reduces not only the computational burden of RNA-seq analysis but also the interpretive discontinuity that often occurs between statistical output and biological insight.

This connected workflow is particularly useful for complex plant transcriptome datasets, where biological questions often depend on expression dynamics across time, genotype, treatment, or environmental condition. In such datasets, the central question is frequently not limited to whether a gene is differentially expressed, but when it changes, in which biological context it changes, and whether it belongs to a functionally coherent gene group. By enabling users to move iteratively between differential expression results, clustering, co-expression modules, gene-set comparison, and functional enrichment, VizR supports a mode of analysis that is well suited to multidimensional plant RNA-seq experiments.

The equalizer-style pattern search extends this framework by enabling hypothesis-guided gene discovery. In contrast to clustering-based approaches, which identify expression patterns after unsupervised grouping, the equalizer interface allows users to define an expected expression trajectory and search the transcriptome for genes that match that pattern. This feature is particularly useful when prior phenotypic or physiological observations suggest the timing or direction of a biological process. In the wound-response dataset analyzed here, this approach recovered late-induced genes associated with secondary barrier formation, illustrating how VizR can translate a user-defined biological trajectory into a directly interpretable gene set.

VizR currently remains optimized for reference genome-based short-read RNA-seq analysis and uses edgeR as its primary differential expression framework. Equalizer-derived outputs should also be regarded as prioritized candidate gene sets that require statistical and experimental validation. In addition, the current implementation is configured primarily for *Arabidopsis thaliana*, and expansion to additional plant reference genomes will further broaden its applicability to diverse plant systems. Nevertheless, by coupling reproducible upstream processing with interactive visualization, reusable gene-set routing, enrichment analysis, co-expression analysis, and pattern-guided gene discovery, VizR reduces the technical fragmentation of RNA-seq analysis while preserving the analytical depth required for biological interpretation. This integrated framework provides a practical and accessible platform for converting plant RNA-seq datasets into biological insight.

## Acknowledgments

This work was funded by the Suh Kyungbae Foundation (SUHF-19010003), the National Research Foundation of Korea (NRF) grant funded by the Korean government (MSIT, No. RS- 2021-NR060084 and RS-2026-25491168), and the Mid-Career Bridging Program through Seoul National University, Republic of Korea to Y.L. W.-T.J. was supported by the Stadelmann–Lee Scholarship Fund at Seoul National University, Republic of Korea.

## Author contributions

Y. L. and W-T.J. conceived the study and designed the system. D.S. established the foundation of the core analysis pipeline. W-T.J. and H.J. built the system and performed the experiments. Y.L. and W-T.J. wrote the manuscript, and all authors revised and approved the final manuscript.

## Competing interest declaration

The authors declare no competing interests.

## Notes

### Competing Interest Statement

The authors have declared no competing interest.

## References

Afgan E, Baker D, Batut B, Van Den Beek M, Bouvier D, Čech M, Chilton J, Clements D, Coraor N, Grüning BA (2018) The Galaxy platform for accessible, reproducible and collaborative biomedical analyses: 2018 update. Nucleic acids research 46: W537–W544

Andrews S (2017) FastQC: a quality control tool for high throughput sequence data. 2010. In, Bolger AM, Lohse M, Usadel B (2014) Trimmomatic: a flexible trimmer for Illumina sequence data. Bioinformatics 30: 2114–2120

Conesa A, Madrigal P, Tarazona S, Gomez-Cabrero D, Cervera A, McPherson A, Szcześniak MW, Gaffney DJ, Elo LL, Zhang X (2016) A survey of best practices for RNA-seq data analysis. Genome biology 17: 13

Danecek P, Bonfield JK, Liddle J, Marshall J, Ohan V, Pollard MO, Whitwham A, Keane T, McCarthy SA, Davies RM (2021) Twelve years of SAMtools and BCFtools. Gigascience 10: giab008

Ge SX, Son EW, Yao R (2018) iDEP: an integrated web application for differential expression and pathway analysis of RNA-Seq data. BMC bioinformatics 19: 534

Haas BJ, Papanicolaou A, Yassour M, Grabherr M, Blood PD, Bowden J, Couger MB, Eccles D, Li B, Lieber M (2013) De novo transcript sequence reconstruction from RNA-seq using the Trinity platform for reference generation and analysis. Nature protocols 8: 1494–1512

Jeon W-T, Kang J, Lee J-M, Kim K, Cheon A, Lee SS, Han M, Jung H, Je S, Yamaoka Y (2026a) Humidity shapes reproductive development in Arabidopsis thaliana by modulating epidermal identity and cell wall dynamics. Cell Reports 45

Jeon W-T, Kim J-A, Cheon A, Lee SS, Kang J, Lee J-M, Lee Y (2026b) A hierarchical abscission program regulates reproductive allocation in Prunus× yedoensis and Prunus sargentii. Horticulture Research 13: uhaf317

Kim D, Paggi JM, Park C, Bennett C, Salzberg SL (2019) Graph-based genome alignment and genotyping with HISAT2 and HISAT-genotype. Nature biotechnology 37: 907–915

Kolberg L, Raudvere U, Kuzmin I, Adler P, Vilo J, Peterson H (2023) g: Profiler—interoperable web service for functional enrichment analysis and gene identifier mapping (2023 update). Nucleic acids research 51: W207–W212

Kolde R, Kolde MR (2015) Package ‘pheatmap’. R package 1: 790

Lamesch P, Berardini TZ, Li D, Swarbreck D, Wilks C, Sasidharan R, Muller R, Dreher K, Alexander DL, Garcia-Hernandez M (2012) The Arabidopsis Information Resource (TAIR): improved gene annotation and new tools. Nucleic acids research 40: D1202– D1210

Langfelder P, Horvath S (2008) WGCNA: an R package for weighted correlation network analysis. BMC bioinformatics 9: 559

Langmead B, Salzberg SL (2012) Fast gapped-read alignment with Bowtie 2. Nature methods 9: 357–359

Lee J-M, Jeon W-T, Han M, Choi M-S, Kwon M, Kim K, Je S, Jung H, Heo G, Joo Y (2025) Wounding induces multilayered barrier formation in mature leaves via phytohormone signalling and ATML1-mediated epidermal specification. Nature Plants 11: 1298–1315

Leinonen R, Sugawara H, Shumway M, Collaboration INSD (2010) The sequence read archive. Nucleic acids research 39: D19–D21

Pertea M, Pertea GM, Antonescu CM, Chang T-C, Mendell JT, Salzberg SL (2015) StringTie enables improved reconstruction of a transcriptome from RNA-seq reads. Nature biotechnology 33: 290–295

Pola-Sánchez E, Hernández-Martínez KM, Pérez-Estrada R, Sélem-Mójica N, Simpson J, Abraham-Juárez MJ, Herrera-Estrella A, Villalobos-Escobedo JM (2024) RNA-Seq data analysis: A practical guide for model and non-model organisms. Current Protocols 4: e1054

Robinson M, McCarthy D, Smyth GK (2010) edgeR: differential expression analysis of digital gene expression data. Bioinformatics 26: 139–140

Schmieder R, Edwards R (2011) Quality control and preprocessing of metagenomic datasets. Bioinformatics 27: 863–864

Shin Y-S, Jeon W-T, Jung H, Do THT, Lee Y (2025) An RbohC and RbohE-mediated ROS oscillatory circuit regulates root cap turnover in Arabidopsis. Iscience 28

Tsai H-H, Tang Y, Jiang L, Xu X, Dénervaud Tendon V, Pang J, Jia Y, Wippel K, Vacheron J, Keel C (2025) Localized glutamine leakage drives the spatial structure of root microbial colonization. Science 390: eadu4235

Upton RN, Correr FH, Lile J, Reynolds GL, Falaschi K, Cook JP, Lachowiec J (2023) Design, execution, and interpretation of plant RNA-seq analyses. Frontiers in plant science 14: 1135455

Wang Z, Gerstein M, Snyder M (2009) RNA-Seq: a revolutionary tool for transcriptomics. Nature reviews genetics 10: 57–63

